# Decoding digits and dice with Magnetoencephalography: Evidence for a shared representation of magnitude

**DOI:** 10.1101/249342

**Authors:** A. Lina Teichmann, Tijl Grootswagers, Thomas Carlson, Anina N. Rich

## Abstract

Numerical format describes the way magnitude is conveyed, for example as a digit (‘3’) or Roman Numeral (‘III’). In the field of numerical cognition, there is an ongoing debate of whether magnitude representation is independent of numerical format. Here, we examine the time course of magnitude processing when using different symbolic formats. We presented participants with a series of digits and dice patterns corresponding to the magnitudes of 1 to 6 while performing a 1-back task on magnitude. Magnetoencephalography (MEG) offers an opportunity to record brain activity with high temporal resolution. Multivariate Pattern Analysis (MVPA) applied to MEG data allows us to draw conclusions about brain activation patterns underlying information processing over time. The results show that we can cross-decode magnitude when training the classifier on magnitude presented in one symbolic format and testing the classifier on the other symbolic format. This suggests similar representation of these numerical symbols. Additionally, results from a time-generalisation analysis show that digits were accessed slightly earlier than dice, demonstrating temporal asynchronies in their shared representation of magnitude. Together, our methods allow a distinction between format-specific signals and format-independent representations of magnitude showing evidence that there is a shared representation of magnitude accessed via different symbols.

## Introduction

Numbers are vital in our everyday life: we need them to count, calculate, and compare. Symbolic notations of numbers allow us to understand and interact with *distinct* quantities. We use a variety of symbolic notations that can all convey the same quantity. For example, the same magnitudes can be expressed using digits (“3”), Roman numerals (“III”), or words (“three”). A central debate in the field of numerical cognition is whether there is a shared brain representation of magnitude or whether representation varies depending on numerical format (Cohen Kadosh & Walsh, 2009).

How does the brain represent magnitude information across different *symbolic* notations?^1^ Most previous studies examining magnitude processes accessed via different symbols, such as digits and number words, have used functional magnetic resonance imaging (fMRI) to compare spatial overlaps of activity (e.g., Eger, Sterzer, Russ, Giraud, & Kleinschmidt, 2003; Naccache & Dehaene, 2001; Pinel, Dehaene, Rivière, & LeBihan, 2001). Although there is some debate about whether numerical processing is independent of notation, the majority of these studies suggest that the intraparietal sulcus (IPS) is critically involved in numerical processing independent of notation type (for reviews see Dehaene, Piazza, Pinel, & Cohen, 2003; Nieder & Dehaene, 2009). While many of these studies show evidence for spatial overlap in the brain’s representation of magnitude across symbols, the dynamic emerging representation of magnitude potentially might have different timing profiles across formats.

Studies using electroencephalography (EEG) have shown that magnitudes presented in different formats are processed similarly over time (Dehaene, 1996; Temple and Posner, 1998; Libertus, Woldorff, and Brannon, 2007). These studies have used univariate analyses to examine magnitude processing over time, averaging activity over many trials to find global activation differences between different stimuli in single EEG channels. A more sensitive approach is multivariate pattern analysis (MVPA), which allows comparison of activity *patterns* (Carlson, Schrater, & He, 2003; Cox & Savoy, 2003; Edelman, Grill-Spector, Kushnir, & Malach, 1998; Haxby et al., 2001; Kamitani & Tong, 2005; Kriegeskorte, Goebel, & Bandettini, 2006; Tong & Pratte, 2012). This approach can test the representational overlap between different symbolic formats of magnitude and, with MEG, how it unfolds over time (Raizada, Tsao, Liu, & Kuhl, 2009). The current study uses MVPA for the time-series neural data (Grootswagers, Wardle, & Carlson, 2016), a novel approach for the field of numerical cognition. We use MEG, which has high temporal resolution, to investigate the timecourse of processing magnitude when accessed via two different symbolic formats: digits and dice. Applying MVPA to time-series neural data allows us to answer the following questions: (1) Is magnitude information conveyed by different symbols (digits and dice) processed in a similar way over time? and (2) can a classifier trained on one numerical symbol successfully generalise to another symbol? Such a finding would be strong evidence in favour of a shared representation of magnitude regardless of notation.

An inherent challenge in studying magnitude processing is the control for visual confounds, because there are unavoidable differences in stimuli representing different magnitudes (e.g., Bulthé et al., 2014; Eger et al., 2009). In our design, we aimed to address this challenge in three ways. First, we presented stimuli in different locations on the screen to add variability to the low-level signals and minimise retinotopic differences between stimuli. Second, we modelled the effects of low-level features to quantify inevitable low-level stimulus differences which could then be regressed out from the magnitude analysis. Third, we drew all of our main conclusions concerning magnitude based on *similarities* of processing magnitude when accessed via two different symbolic notations: digits and dice. As the low-level features of dice do not vary in the same way as those of digits, the key results cannot be driven by low-level features. Using these careful controls to minimise the effects of visual feature differences, we addressed the key question of whether there is a shared representation of magnitude across symbolic notations.

## Methods

### Participants

Twenty participants (14 female, mean age = 28.5 years, SD = 8.6, age range: 20 – 51 years) completed the study. All participants reported normal or corrected-to-normal vision. Participants gave informed consent before the experiment and were reimbursed with $20/hour. During MEG recording, participants were asked to complete a magnitude 1-back task (see below) to ensure they attended the stimuli. One participant performed more than two standard deviations below the group mean in this task and was therefore excluded from analysis, leaving 19 participants in total (13 female, mean age = 28.5 years, SD = 8.8; age range: 20 – 51 years). The study was approved by the Macquarie University Human Research Ethics Committee.

### Procedure

Participants completed 8 blocks of a 1-back task (Figure 1) while lying in a dimly lit magnetically shielded room (MSR) for MEG recordings. Each block contained 216 trials. In each trial, participants were presented with a black fixation cross and four black outlined squares as placeholders around it. The presentation duration of the fixation screen varied on a trial-to-trial basis between 900 and 1200ms. Then, a black numerical symbol appeared in one of the four placeholders while the fixation cross and four squares remained visible. The squares surrounding each stimulus were at 2.85 degrees visual angle. The horizontal and vertical distances between these squares were at 6.9 and 8.8 degrees visual angle, respectively. We used two different numerical symbols as format (dice or digits) with magnitudes 1 to 6. Overall there were 48 different stimuli (4 locations, 2 formats, 6 magnitudes) which were repeated 32 times throughout the experiment. Stimuli remained on the screen for 83ms. Participants were asked to push a button if the same magnitude repeated, regardless of location (four squares) or numerical format (digits or dice). There were 24 such repeat-trials per block in which participants were meant to press the button. These trials were excluded from analysis. Response time was limited to a maximum of 800ms after stimulus onset. Participants received feedback on their accuracy after each block. Participants were instructed to fixate on the fixation cross throughout the experiment.

**Figure 1:**
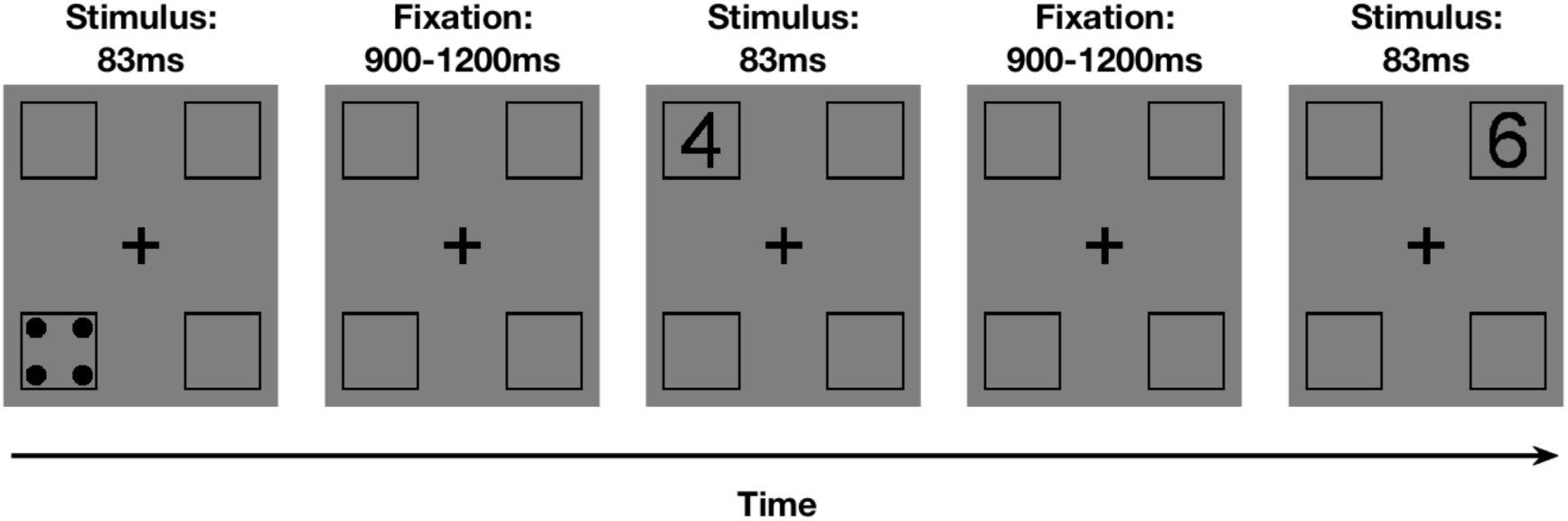
On every trial participants were presented with a magnitude between 1 and 6 in one of two different numerical symbols (digits or dice) in one of four locations. The possible locations were framed in black. Then a fixation screen was presented for a variable duration between 900 and 1200ms. The fixation duration was sampled at random from a uniform distribution. The task was to press a button when the same magnitude repeated on consecutive trials. During the post-stimulus fixation period, participants had a maximum of 800ms to respond.

### Apparatus and Pre-processing

Before the MEG recordings, participants’ head shapes were measured using a digitiser pen (Polhemus Fastrak, Colchester, USA). We used the digitiser pen to register three reference locations (left and right preauricular and nasion) and the locations of five marker coils. Participants wore an elastic cap with the marker coils throughout the session to measure the head position before and after the experiment. During the MEG recording, stimuli were projected onto a translucent screen mounted in the MSR. MATLAB with Psychtoolbox extension (Brainard, 1997; Kleiner et al., 2007; Pelli, 1997) was used for stimulus presentation. Button presses were recorded with a Bimanual 4-Button Fiber Optic Response Pad (Current Designs, Philadelphia, USA). Participants held one of the response pads in their hands and were instructed to press the button with their thumb. The neuromagnetic recordings were obtained with a whole-head axial gradiometer MEG (KIT, Kanazawa, Japan). The system has 160 axial gradiometers and recorded at 1000Hz. An online low-pass filter of 200Hz and a high-pass filter of 0.03Hz was used. We determined stimulus onsets with a photodiode that detected light change when a number stimulus came on the screen. We used FieldTrip (Oostenveld, Fries, Maris, & Schoffelen, 2011) for all preprocessing steps. Trials were epoched from ‐100 to 800ms relative to the onset of the stimulus and downsampled to 200Hz (5ms resolution). Next, to the reduce dimensionality of the data, we used principal component analysis (PCA) and retained the principal components that explained 99% of the variance in the data for each participant. Following a standard analysis pipeline by Grootswagers et al., (2016), we performed no further pre-processing steps (e.g., channel selection, artefact correction). This maintains the data in the rawest possible form.

### Pattern Classification

We used both a decoding analysis approach and representational similarity analysis (RSA, Kriegeskorte, 2011; Kriegeskorte & Kievit, 2013; Kriegeskorte, Mur, & Bandettini, 2008) to decode magnitude over time. In the following, we address each approach in turn.

### Decoding Analysis

For a decoding analysis, patterns of brain activity for each participant are extracted across all MEG channels (components after PCA). A linear discriminant classifier is trained to distinguish between brain activity patterns evoked by all stimuli. Then an independent subset of data from the same participant is used to test whether the classifier can predict which stimulus evoked a certain pattern of activity. The training and testing steps are repeated at every time-point. If the prediction is above chance at a given time we can infer that the information the classifier had in the training phase is relevant for the prediction at that time-point.

We used random-effect Monte-Carlo cluster statistics corrected for multiple comparisons (as implemented by CosmoMVPA toolbox, (Oosterhof, Connolly, & Haxby, 2016; Maris & Oostenveld, 2007) to determine whether the classifier performed above chance. Threshold Free Cluster Enhancement (TFCE, Smith & Nichols, 2009) was used as a cluster-forming statistic. The TFCE statistic represents the support from neighbouring time points, allowing optimal detection of sharp peaks, as well as sustained weaker effects. To correct for multiple comparisons, the Monte-Carlo technique used by CosmoMVPA performs a sign-permutation test, swapping the signs of the decoding results of all participants at random at each time point and recomputes the TFCE statistic. This is repeated 10,000 times to obtain a null distribution at each time point. Then the most extreme value of each null distribution is taken in order to construct an overall null distribution across the time-series. The 95^th^ percentile of this overall null distribution is used when we compare the real decoding results and the null hypothesis providing a p-value (*α* = 0.05) which is corrected for multiple comparisons.

We ran two decoding analyses: within-format and between-format classification. For the within-format classification, we trained a classifier on magnitude (values 1-6) using a subset of digit trials and then tested the classifier on digit trials of independent data (chance level = 16.67%). We repeated the same process for the dice stimuli. In the 32-fold crossvalidation, each fold contained data corresponding to 24 individual stimuli: Each magnitude (1-6) was repeated 4 times, once in each of the four locations (top right, top left, bottom right, bottom left). Each of the folds served as independent test data once while all other folds were used for classifier training. For the between-format analysis, we trained the classifier on magnitude (values 1-6) in all digit trials and then tested its performance on data from the dice trials and *vice versa*.

It is important to note that the within-format classifiers can make use of magnitude *and* visual information to predict which magnitude evoked a given pattern of brain activity. To decrease the contribution of low-level visual differences we presented the stimuli in four different locations and hence reduced retinal overlap. While this approach increases the variability in the stimuli there is still a considerable degree of low-level feature similarity in the stimuli (e.g., total density, edges, orientation, curves). That means we cannot draw definite conclusions about magnitude processing from the within-format analysis. In comparison, the classifier in the between-format analysis was trained on magnitude in one notation (e.g., digits) and tested on the other notation (e.g., dice). That means the between-format classifier can only rely on magnitude information, making this the strongest test of the hypothesis that there is a representation of magnitude that does not depend on the specific symbol of presentation.

### Representational Similarity Analysis (RSA)

Using RSA (Kriegeskorte, 2011; Kriegeskorte & Kievit, 2013; Kriegeskorte, Mur, & Bandettini, 2008), we quantified the similarity between brain activity patterns evoked by different stimuli. First, we averaged the trials corresponding to the 48 unique trials (i.e., unique combinations of format, location, magnitude) and correlated these average trials with one another. High correlations indicate that the evoked activity is similar for a given pair of stimuli and therefore harder to distinguish. We correlated all possible stimulus pairs at each time-point and thus ended up with a total of 180 representational dissimilarity matrices (RDMs, see Figure 2). We then constructed five different model RDMs, two conceptually based (Magnitude Model, Label Model) and three visually-based models (Location Model, Silhouette Model, and Format Model). We tested whether these models could capture the differences in the neural MEG RDMs by correlating these models to the neural RDMs. In the same way as for the decoding analysis, significance was tested with the Monte-Carlo cluster statistics corrected for multiple comparisons.

**Figure 2:**
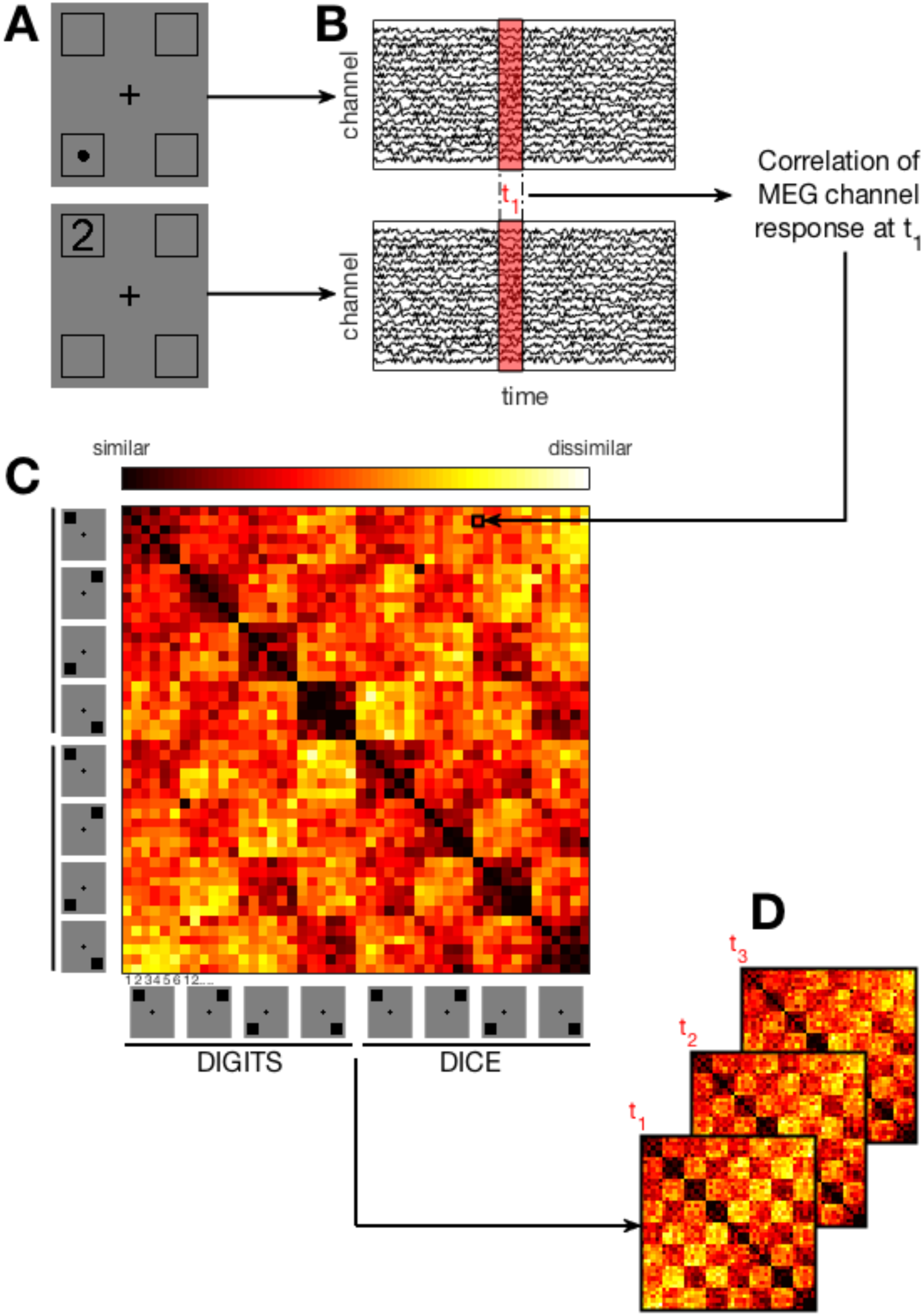
Panel A depicts stimuli seen in two separate trials. Panel B shows the recorded MEG signal in response to these stimuli. The signals from both trials are then correlated at each timing window (e.g., t_1_). The correlation values of each stimulus pair are then inserted into the dissimilarity matrix of the corresponding timing window (Panel C). This process is repeated for all stimulus pairs and every time window to create a time series of dissimilarity matrices (Panel D).

Our key model was the Magnitude Model (Figure 3A). The Magnitude Model is based on the theory that magnitudes are represented on a mental number line (Moyer & Landauer, 1967; Restle, 1970). The Magnitude Model hence predicts that correlations of stimulus pairs that are closer together in magnitude (e.g., 1 and 2) will be higher than correlations of stimuli that are farther apart (e.g., 1 and 5). In the Magnitude Model, location and format are irrelevant, the prediction depends solely on magnitude.

**Figure 3:**
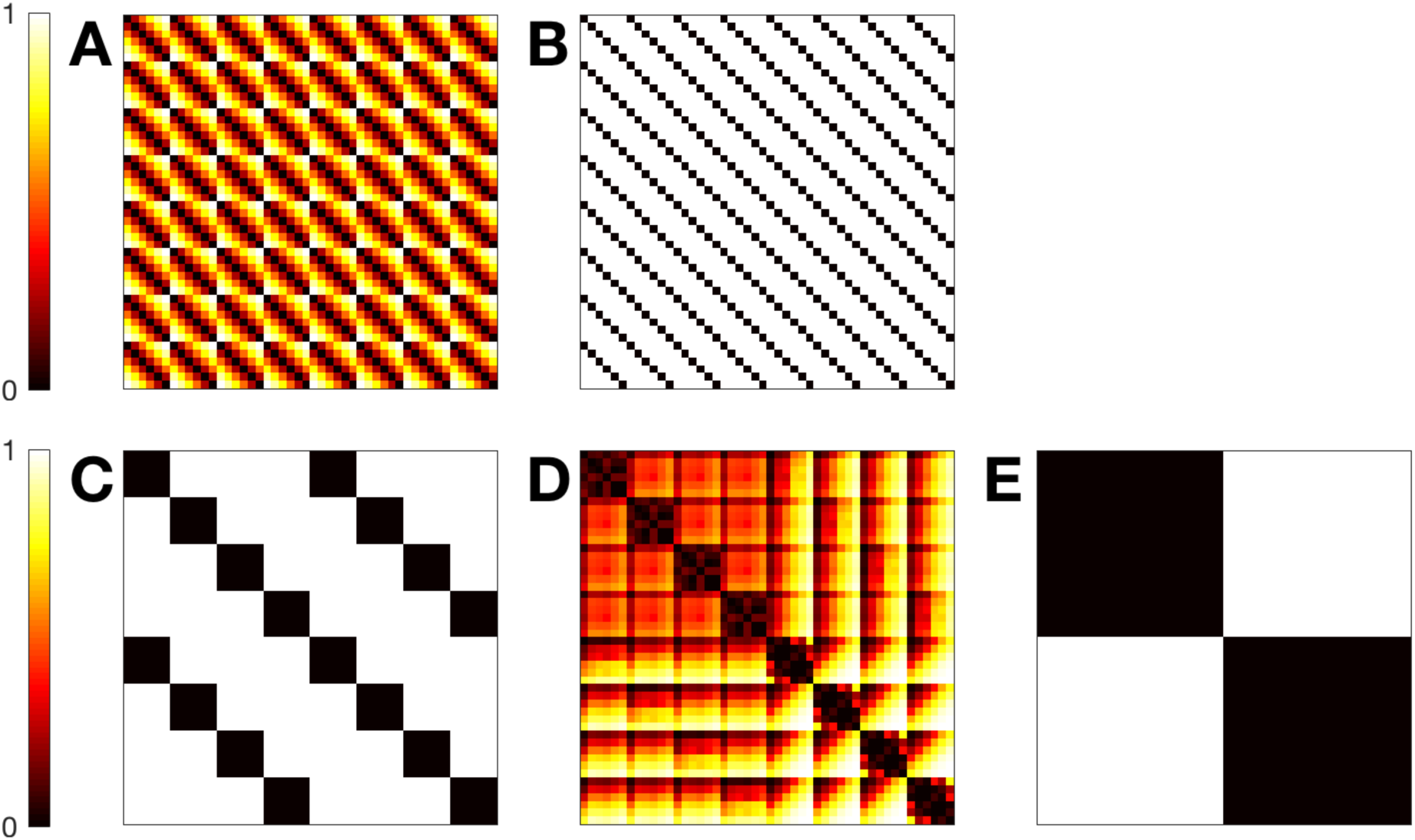
The model Representational Dissimilarity Matrices (scaled). The top row shows the conceptually based Magnitude and Label Models (Panels A and B, respectively). The bottom row shows the visually-based Location, Silhouette, and Format Model (Panels C-E). Each square represents the predicted dissimilarity between a stimulus pair where 1= highly dissimilar and 0 = highly similar.

The Label Model (Figure 3B) served as a control model. As participants were required to detect when a magnitude repeats, it is plausible that correlations of the neural MEG RDMs and the Magnitude Model RDM could be driven by a verbal labelling strategy of participants. The Label Model coded for such strategy by predicting that data evoked by stimuli with the same verbal labels (e.g., 1 presented in both numerical formats) would have a high correlation while stimuli with different verbal labels (e.g., 1 presented as a digit in the top left and 2 presented as a dice in the bottom right) would have a low correlation. This model assumes that all number pairs that do not have the same verbal label are equally hard to distinguish.

We constructed the visually-based models to examine what part of the correlations between the MEG RDMs can be explained by inevitable low-level visual differences between the stimuli. The Location Model (Figure 3C) models in which of the four squares the stimulus was presented. The Location model ignores magnitude and format and predicts that correlations of stimuli presented in the same location are higher than correlations of stimuli which were presented in different locations. The Silhouette Model (Figure 3D) compares visual overlap between the stimuli (Jaccard, 1901). The prediction for the visual model is that the brain activity pattern evoked by stimuli which have more pixel overlap also have a higher correlation than patterns evoked by stimuli that do not have as much visual overlap. Lastly, the Format Model (Figure 3E) predicts that data of trials in which the numerical format is the same (e.g., digits and digits) will correlate stronger than data of trials with different numerical format (e.g., digits and dice). The Format Model ignores location and magnitude and solely codes for format.

## Results

In the 1-back task, participants accurately detected 82.2% (SD = 8.3%) of the repeat-trials. To analyse the MEG data, we ran a decoding analysis and RSA. We will first present the results from the decoding analysis and then the results from the RSA.

### Decoding Analysis

For the within-format decoding, the classifier was trained and tested on stimuli of the same numerical format and hence can be driven by both visual and magnitude information. The classifier was able to predict the numerical value above chance for a cluster stretching from 120 to 740ms relative to stimulus onset for dice and from 145 to 475ms for digits. The within-dice classifier performance is above chance for a longer period of time in comparison to the within-digit classifier performance (Figure 4), presumably reflecting the stronger visual differences present in the dice stimuli from 1-6. This means that when the classifier is trained on magnitudes of the same numerical format, it is able to distinguish the classes above chance for a substantial period of the time series. Even though stimuli were presented in different locations (right/left, bottom/top), visual features such as comparisons). Under the graph are the sensor contributions (arbitrary units) to the decoding results. shape will contribute to classifier performance. This is in line with the finding that classifier performance for dice is more accurate than for digits: dice have a more distinct visual pattern than digits, and more visual information corresponds to a higher magnitude value, confounding possible coding of magnitude with visual differences.

**Figure 4:**
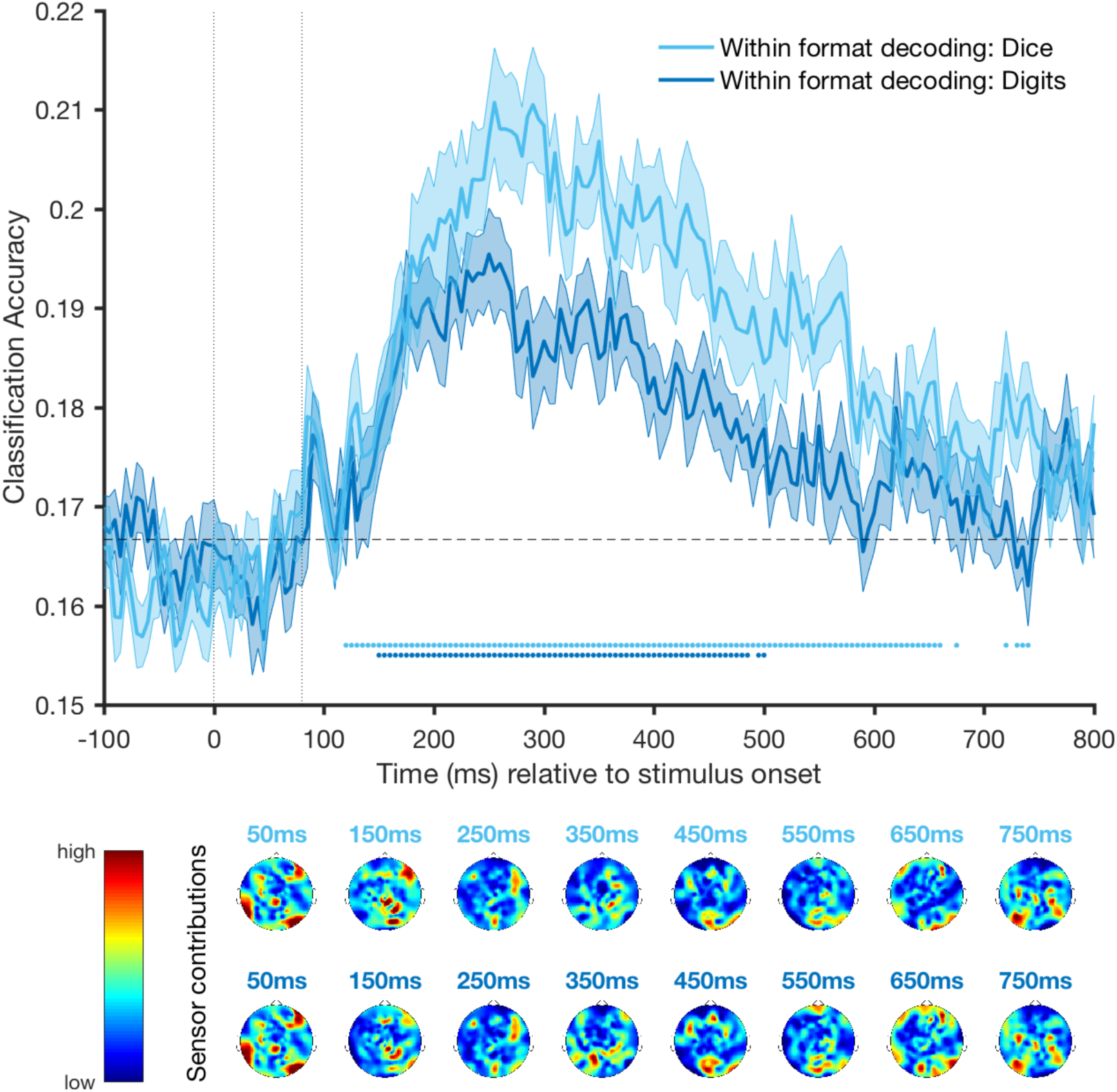
Classification accuracy over time for within-format decoding of dice (light blue) and digits (dark blue). Shading indicates standard error around the mean. The dashed horizontal line shows chance level while the dotted vertical lines show the stimulus duration. The coloured dots indicate classification accuracy that is significantly above chance (p<0.05, corrected for multiple

In the between-format decoding, we trained a linear discriminant classifier on the magnitude (values 1-6) of one format (e.g., dice) and tested its performance on the other format (e.g., dice) and *vice versa.* In comparison to the within-format decoding, there are no reliable visual differences between stimuli in the between-format decoding analysis that could predict above chance classification, making this a strong test of the hypothesis that a shared representation of magnitude exists. The results for the between-format decoding (Figure 5) show that there is cluster of classifier performance above chance stretching from 410 to 435ms when the classifier was trained on dice and tested on digits. When the classifier was trained on digits and tested on dice, it performs significantly above chance in a cluster between 390 and 485ms. As low-level features such as density do not vary systematically for digits, classification is most likely driven by magnitude demonstrating a shared representation of magnitude accessed via digits and dice.

**Figure 5:**
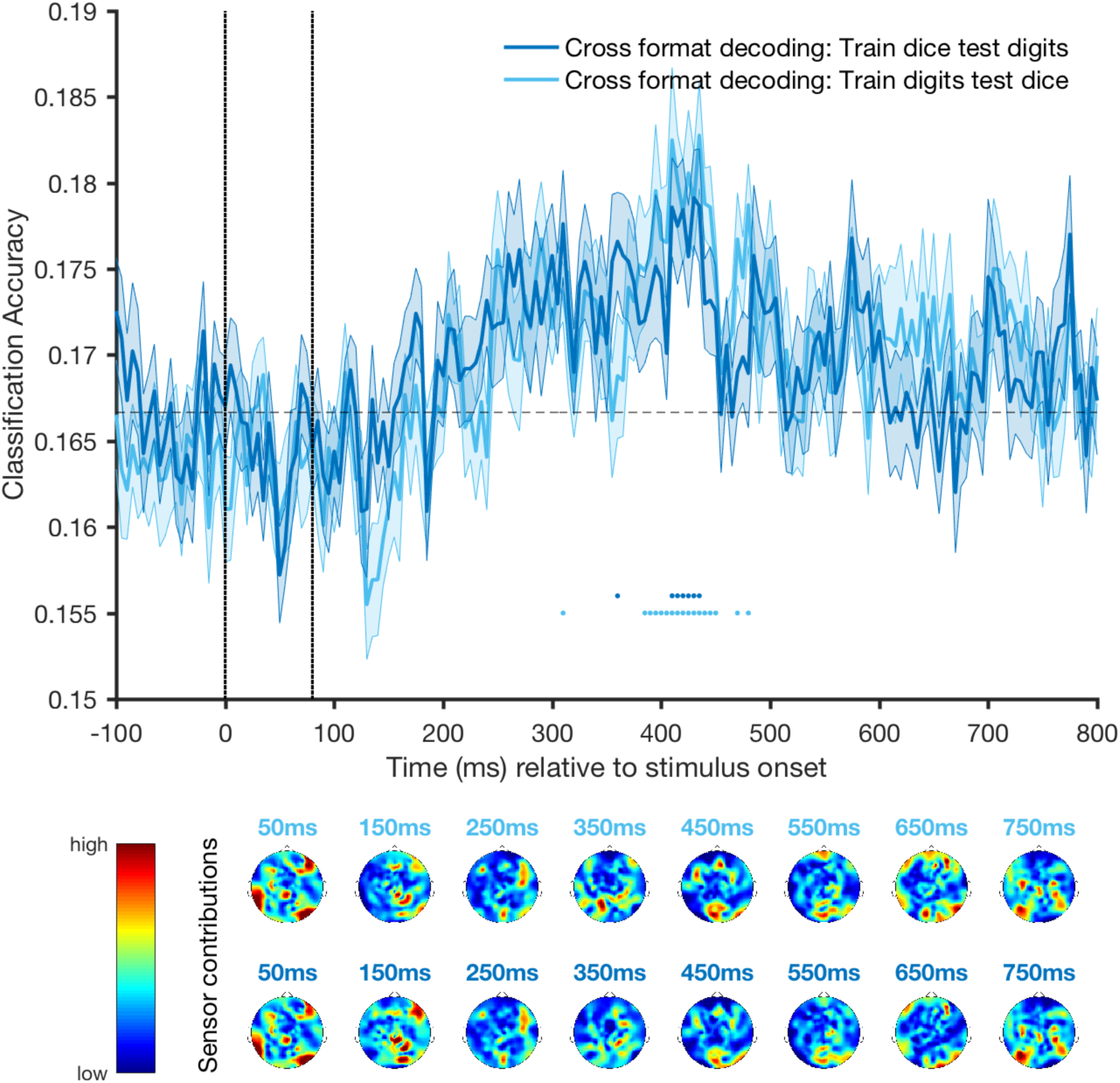
Classification accuracy over time for between-format magnitude decoding when the classifier is trained on dice and tested on digits (light blue) and vice versa (dark blue). Shading indicates standard error around the mean. The dashed horizontal line shows chance level while the dotted vertical lines show the stimulus duration. The coloured dots indicate classification accuracy that is significantly above chance (p<0.05, corrected for multiple comparisons). Under the graph are the sensor contributions (arbitrary units) to the decoding results.

The significant between-format classification suggests that there is overlap in the representation of digits and dice. We now compare the relative time it takes to access magnitude information from the two formats using a time-generalisation technique. It is, for example, possible that one format is processed faster than the other one and we have only captured a slight overlap between their processing time-windows with the between-format decoding. To test this possibility we examined whether the between-format decoding generalises over time (Carlson, Hogendoorn, Kanai, Mesik, & Turret, 2011; King & Dehaene, 2014). We trained the classifier on trials of one numerical format (e.g., digits) at each time point of the time-series and then tested the classifier on the other numerical format (e.g., dice) at every possible time point (Figure 6A). To test this difference statistically, we conducted a random-effect Monte-Carlo statistic that is corrected for multiple comparisons to find which time points in the time-generalisation matrix have classification that is above chance. This allows us to see whether a brain activity pattern that was observed for digits at a given time point appeared in a similar way for dice at a later or earlier time point (or *vice versa*). The results for the time-generalisation analysis are summarised in Figure 6. The red line (Figure 6B) indicates the expected between-format decoding if training and testing time correspond perfectly (no temporal asynchrony in processing digits and dice). However, visual inspection suggests that, relative to this diagonal, there is a rightward shift when we train on dice and test on digits, and a leftward shift when we train on digits and test on dice. We then calculated the distance between the significant time points to the red diagonal reference line that indicates a perfect one-to-one temporal mapping. The results show that the time generalisation of the classifier performance is shifted later by a median of 40ms when we trained on dice and tested on digits, and 45ms earlier when we trained on digits and tested on dice (Figure 6C). This shows that there is indeed a time shift between the processing speed of magnitudes presented as digits and dice: When training the classifier on dice we are able to generalise to digits earlier and *vice versa,* suggesting that access to magnitude information occurs earlier for digits than dice. However, it is important to note that magnitudes accessed via digits and dice must be similar as the between-format classification is possible. From the decoding analysis, we can hence conclude that there is a representational overlap between accessing magnitude from digits and dice, but that digits appear to be accessed slightly faster than dice.

**Figure 6:**
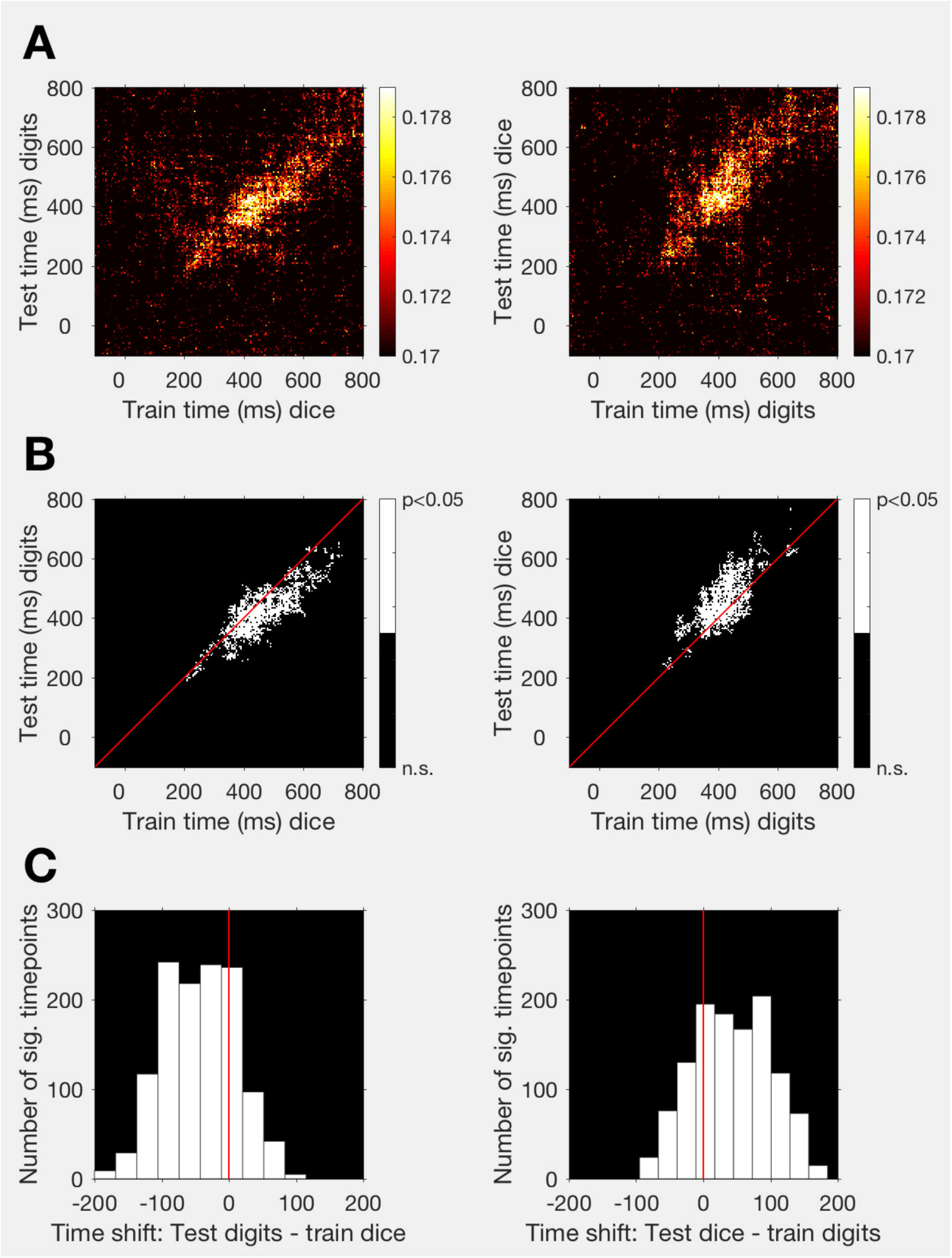
Time generalisation for between-format decoding. Row A shows the classification accuracy across training and testing time when the classifier is trained on dice and tested on digits (left) and vice versa (right). The diagonal line in row B indicates what exact temporal mapping between training and testing time would look like. The white points are train-and test-time combinations where classification is significantly above chance (p<0.05, corrected for multiple comparisons). Row C shows the time-shift from the diagonal of all significant timepoint combinations.

### Representational Similarity Analysis (RSA)

RSA allows us to compare the overall similarity of the brain activity corresponding to *all* of our stimuli instead of only comparing stimulus pairs. We constructed five different models that we compare to the neural MEG RDMs at every time point (Figure 3). This enables us to model what type of information is most prevalent in the signal at a given time. We first correlated the model RDMs with the MEG RDMs at every time point using Spearman’s Rank Correlation. We then used randomeffect Monte-Carlo cluster statistics to quantify whether the correlations were significantly above zero. The results of the RSA are summarised in Figure 7. Stimulus onset and offset are shown by the vertical lines. The black dotted line shows the lower bound of the noise ceiling (Nili et al., 2014), defined as the average correlation between individual subject RDMs and the mean of all other subject RDMs. The noise ceiling is an estimation of how well the true model could perform given the noise in the data (Nili et al., 2014). The noise ceiling highlights that we can expect the maximum correlations between any model RDM and the data to be relatively low just before and after stimulus onset (i.e., more noise in the data) and at the end of the time-series. The noise ceiling peaks at 150ms after stimulus onset indicating that there is less noise in the data at that time point in comparison to earlier or later time points. As a consequence, models that explain the data well in that time frame will have higher correlations compared to models that perform well a little later. This is clearly the case when we look at the correlation between the MEG RDMs and the visually-based models (i.e., Location Model, Visual Model, and Format Model). Visual stimulus characteristics that allow our visual system to distinguish stimuli by features such as shape and location are available relatively early (Ramkumar, Jas, Pannasch, Hari, & Parkkonen, 2013; VanRullen & Thorpe, 2001), leading to high correlations between the visually-based model RDMs and the MEG RDMs early in the time-series. The correlation between the MEG RDMs and the Location Model approaches the noise ceiling early and stays significantly above chance for almost the whole time-series (significant cluster of time points from 55 to 800ms). The Silhouette Model codes for the shape of the stimuli by comparing pixel overlap. The Silhouette Model correlates strongly with the MEG RDMs and peaks at the same time as the Location Model at around 150ms after stimulus onset. The Format Model that codes for whether the magnitude was conveyed by a digit or die correlates significantly above zero with the MEG RDM at a cluster stretching from 145 to 800ms after stimulus onset. The correlation between the Format Model and the MEG RDMs peaks later than the other two visually-based models at 255ms after stimulus onset.

**Figure 7:**
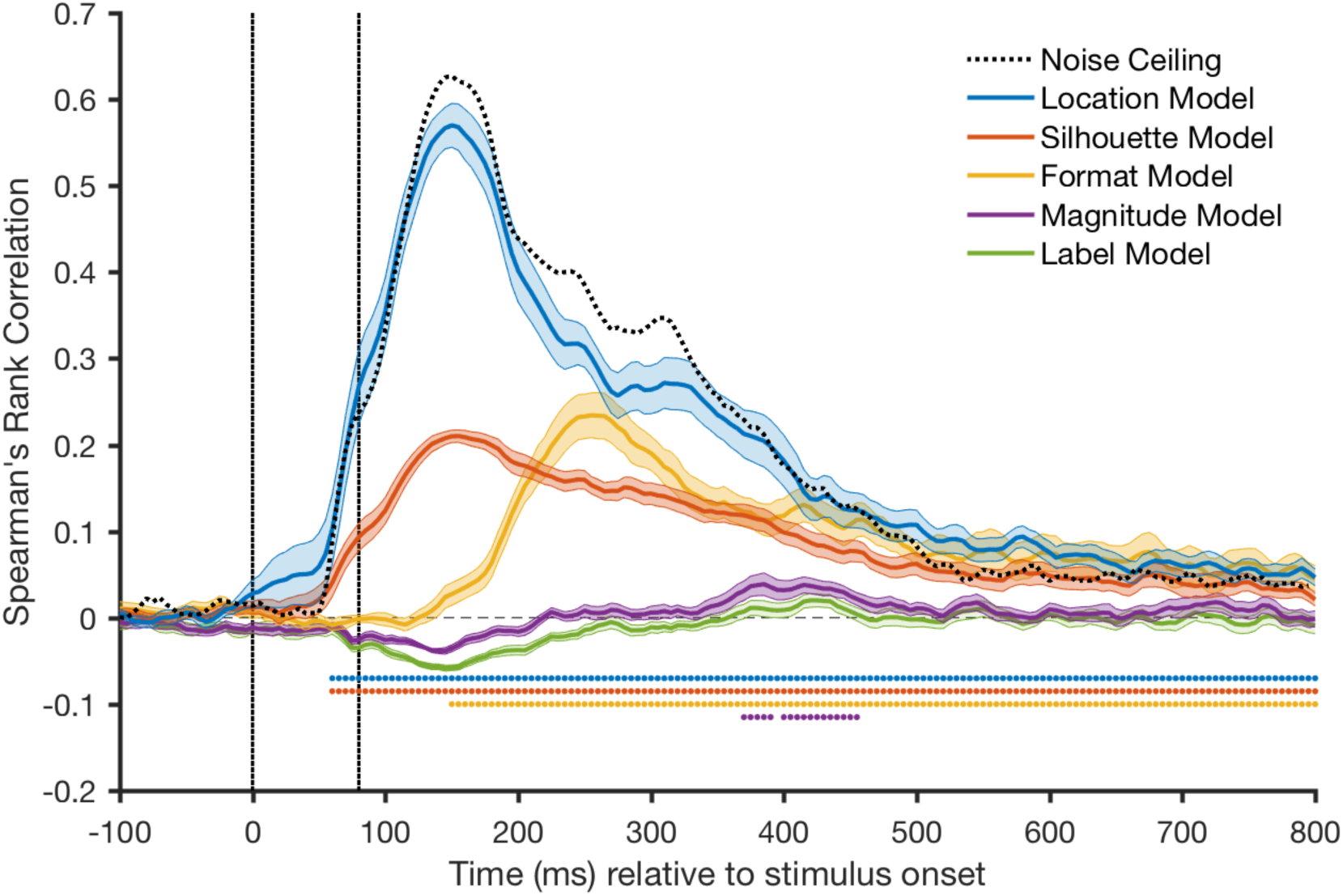
Spearman’s Rank Correlation of all Model Representational Dissimilarity Matrices (RDMs) and the MEG RDM over time. The vertical dotted lines indicate how long the stimulus was on the screen. Each coloured line depicts the correlation of a Model RDM and the MEG RDM. Shades around lines depict standard errors. Coloured dots indicate correlations that are significantly (p<0.05, corrected for multiple comparisons) above zero.

Our key Magnitude Model is an ordinal model predicting that data evoked by stimuli with numerical values that lie closer together (e.g., 1 and 2) should correlate more than data evoked by stimuli with numerical values that are farther apart (e.g., 1 and 6). The Magnitude Model has a correlation larger than zero with the MEG RDMs in a cluster stretching from 365 to 455ms after stimulus onset. This onset corresponds to the significant time windows of the between-format decoding analysis. Note that the correlation between Magnitude Model and MEG data at the significant time points is much lower than for the visually-based models. This also matches the results of the decoding analysis which showed that the mainly visually-driven within-format classification is more accurate than the between-format decoding. There may be several reasons for the absolute difference between visual and magnitude effects. First, as the magnitude effect appears later than the visual effects, the correlation will always be weaker because the data are much noisier by that stage relative to the strong early effects (compare the noise ceiling at these time points; Figure 7). Second, magnitude effects are likely to be more strongly influenced by individual differences than visual effects, as more processing is required to access magnitude than the low-level early sensory signals. Importantly, despite the absolute differences, our results suggest that magnitude is represented independently of location and format.

One possible alternative interpretation of the correlation of the MEG RDMs and the Magnitude Model is that participants internally labelled the stimuli (e.g., “one” regardless of whether dice or digit was presented) to assist with completion of the 1-back task. To test this, we also used a Label Model coding for same versus different verbal label. Although the correlation between the MEG data and the Label Model follows the shape of the Magnitude Model, it does not reach significance at any point throughout the time-series. To test whether the Magnitude Model explains more of the variance than the Label Model, we tested for differences between the correlations of the data with the Magnitude and Label Model. Perhaps not surprisingly given how close the models are to chance, the difference between Magnitude and Label Model did not reach significance at any point in the time-series. Therefore, we cannot rule out a contribution of labelling in the correlation of the Magnitude Model with the MEG data. What we have, however, is evidence that the Magnitude Model explains a significant portion of data variance, in the absence of such evidence for the Label Model (which could reflect insufficient power or an actual lack of an effect).

The initial RSA analysis shows that visual information is strongly correlated with the data but that magnitude information arises in the signal at a later point in the time-series. Looking more closely at the Magnitude Model, we see that in the beginning of the time-series there is a negative correlation between model and MEG data. This negative correlation coincides with the time-point at which the Location and Visual Model peak. That indicates the Location and Silhouette Models account for variance for which the Magnitude Model cannot account. In the next step, we regress out the variance explained by the Location and Silhouette Model and look at the Magnitude Model again (Figure 8). This effectively removes the visual "noise" from the Magnitude Model correlation. Regressing out the Location and Silhouette Model improves the Magnitude Model correlation early in the timeseries. This improvement is more pronounced when the Location Model is removed compared to the Silhouette Model (reflecting the greater correlation between the data and the Location Model compared with the Silhouette Model). Importantly, there is a significant correlation between the Magnitude Model and the MEG data regardless of whether any visual information is regressed out.

**Figure 8:**
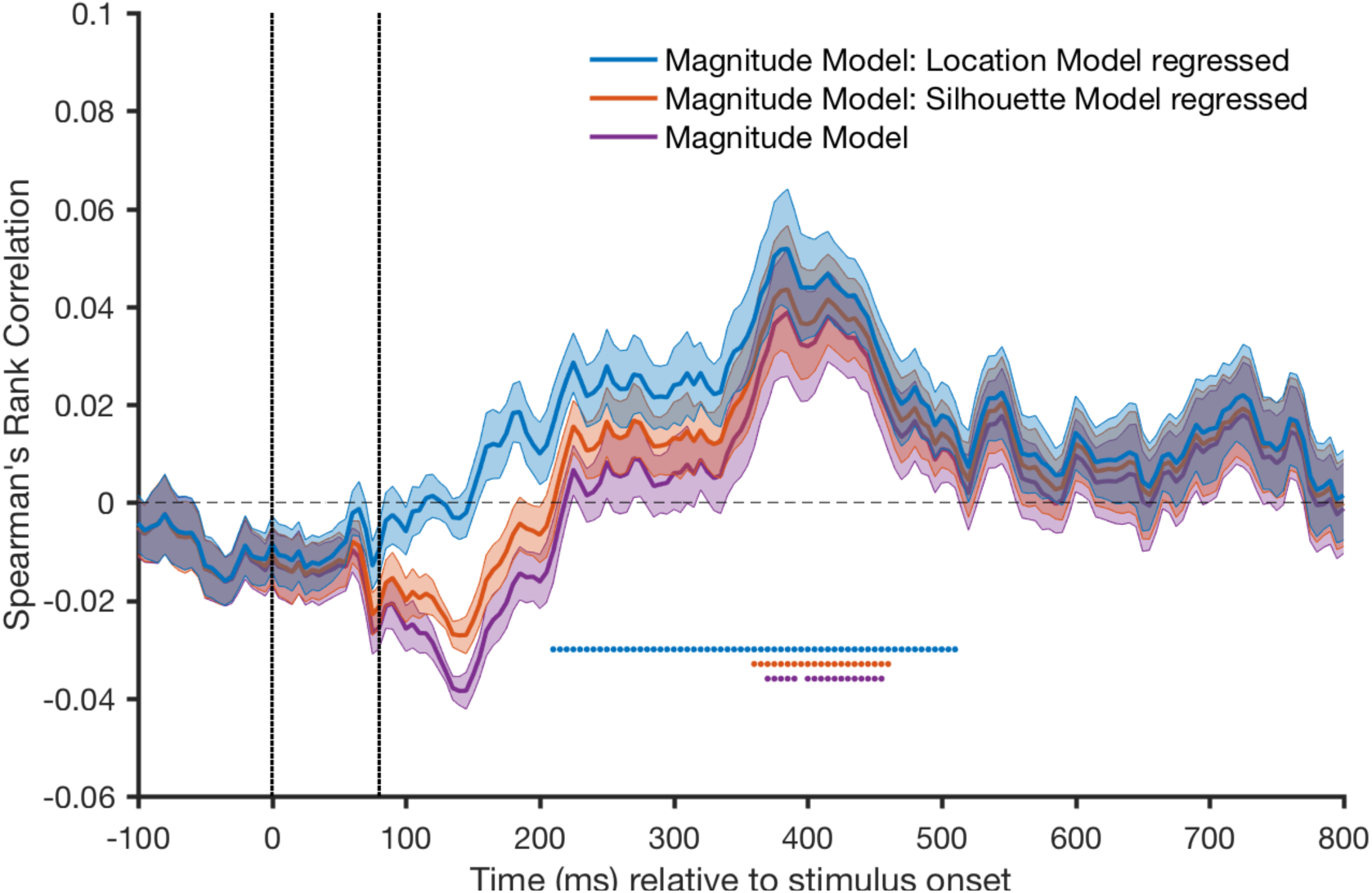
Spearman’s Rank Correlation of the Magnitude Model RDM and the MEG RDM over time when variance explained by the location and silhouette model are regressed out (blue and orange line, respectively) versus when nothing is regressed (purple line). Shading represents standard errors. Coloured dots indicate correlations that are significantly (p<0.05, corrected for multiple comparisons) above 0.

After regressing out the variance accounted for by the Location Model we looked at the Magnitude Model correlation in more detail. The Magnitude Model predicts data evoked by stimuli with numerical values close to one another to be more similar than data evoked by stimuli with numerical values farther apart *independent* of location and format. That means the model contains both within-format and between-format correlations. Before drawing conclusions about the representation of *magnitude* then, it is important to test whether the correlation of the Magnitude Model and the MEG data could be driven by only the within-format correlations, which we know are influenced by visual features. In the next step, we therefore looked at the Magnitude Model separated into three parts: within-digits, within-dice and between-format correlations. The results (Figure 9) show that there is a significant correlation for all three.

**Figure 9:**
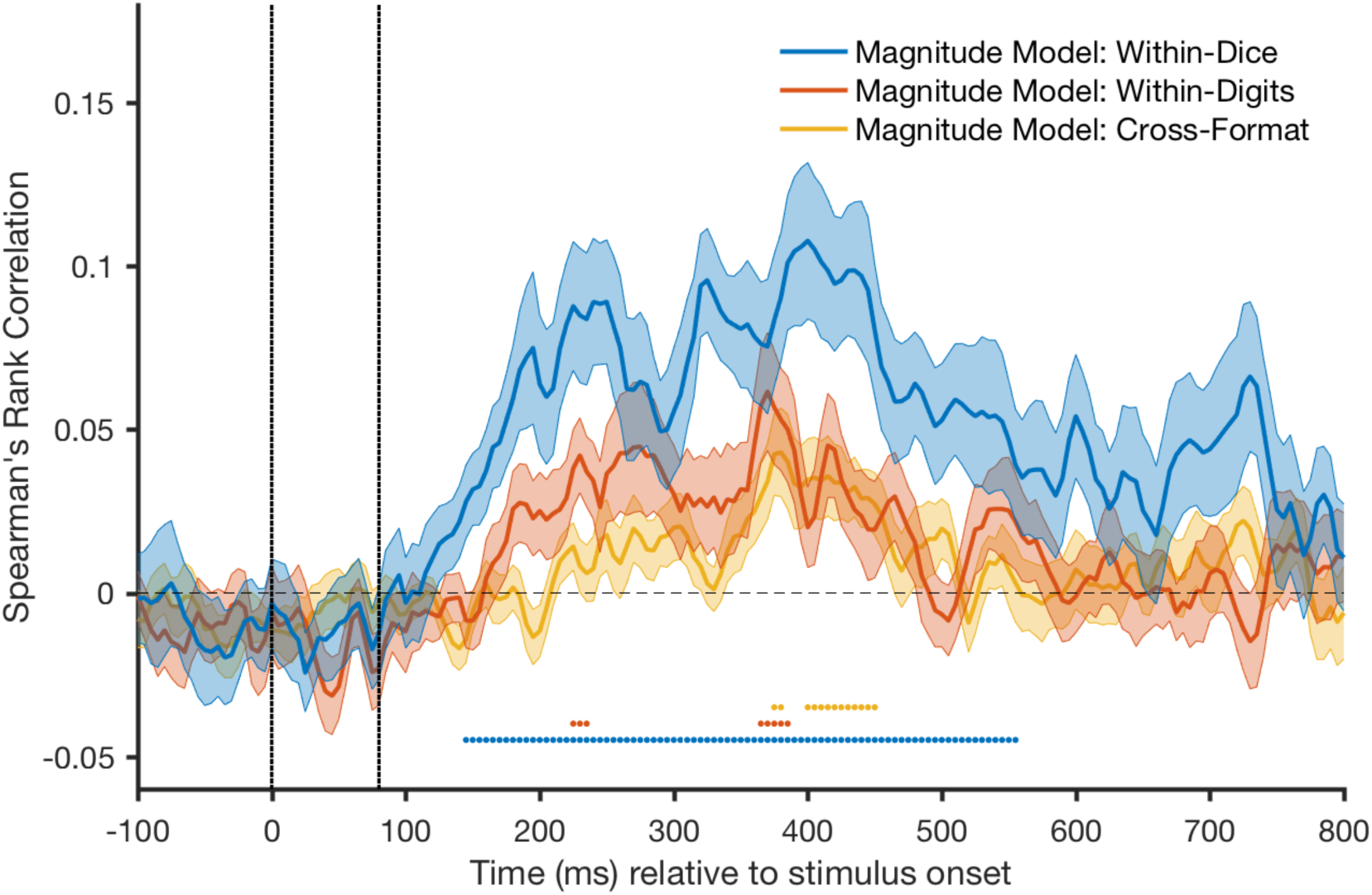
Spearman’s Rank Correlation for different parts of the Magnitude Model Representational Dissimilarity Matrix (RDM) and the MEG RDM when variance explained by the Location Model is regressed out. Shading represents standard errors. Coloured dots indicate correlations that are significantly (p<0.05, corrected for multiple comparisons) above 0.

Supporting the results of the decoding analysis, the within-dice part of the Magnitude Model has the highest correlation with significant clusters stretching over most of time-series (140ms – 735ms relative to stimulus onset). The within-digits part of the Magnitude Model also significantly correlates with the data at several clusters throughout the time-series (225ms – 420ms relative to stimulus onset). Again, it is important to note that most of these correlations are due to visual features and it is not possible to determine any effect of magnitude information alone from these within-format contrasts. In comparison, the between-format part of the Magnitude Model only predicts similarity between data evoked by a certain magnitude in one format (e.g., digit 3) and the same magnitude in the other format (e.g., die 3), thus containing similarities based on magnitude only. This between-format part of the Magnitude Model correlates significantly with the data at a cluster stretching from 360ms to 450ms relative to stimulus onset. This time-window is consistent with the results of the between-format decoding analysis. Thus, these results suggest that magnitude is represented in a similar way when accessed via digits and dice.

## Discussion

In this study, we examine whether there is a common representation of magnitude regardless of symbolic notation (digits and dice). Consistent across two different analysis methods, our results suggest that there is a shared brain representation of magnitude for these symbolic formats. We also see a time difference in the access to this magnitude representation, with digits being processed slightly earlier than dice. In addition, we showed that activation patterns evoked by stimuli closer in numerical value are more similar than of stimuli farther apart, providing neural underpinnings for an ordinal component of magnitude representation.

Previous studies examining magnitude representation have mostly focussed on whether magnitudes presented in different numerical formats are processed in the same brain area (e.g., Eger, Sterzer, Russ, Giraud, & Kleinschmidt, 2003; Naccache & Dehaene, 2001; Pinel, Dehaene, Rivière, & LeBihan, 2001). In the current study, we used a time-series decoding approach to investigate the temporal unfolding of magnitude processing. Our results show that digits and dice are processed in a sufficiently similar way over time to allow for cross-generalisation and that digits and dice which represent closer magnitudes are more similar in neural activity than those that are farther apart. This is in line with behavioural findings such as the numerical distance effect (Moyer & Landauer, 1967) which has been shown to occur independent of numerical format (Schwarz & Ischebeck, 2000).

In addition to similarities in magnitude representation of digit and dice stimuli over time, our results show that there is a temporal shift when comparing the processing of magnitude in these formats. Magnitude from digit stimuli seems to be accessed earlier than for dice. This corresponds to previous behavioural findings by Buckley and Gillman (1974) showing that reaction times to digits are faster than to dots in a regular, known composition. In previous EEG studies, digits have also been shown to be processed slightly earlier than number words (Dehaene, 1996) and dots in random configurations (Temple and Posner, 1998). Similarly, the results of the time-series decoding analysis suggest that magnitude information from digits is accessed slightly earlier than in dice.

Evidence for a similar pattern in processing of magnitude across formats has previously been taken as evidence for *abstract* magnitude representation (see for example Cohen Kadosh & Walsh, 2009). In the context of numerical cognition, “abstract representation” means that magnitude is accessed via a transformation of numerical stimuli to a format-independent, continuous quantity (Dehaene, Dehaene-Lambertz, & Cohen, 1998). This is one possible interpretation of our findings: it may be that digits and dice are both converted into a completely abstract representation of magnitude. The delay between accessing magnitude for dice stimuli in comparison to digits could then be attributed to a difference in conversion speed. It may be that it is faster to access abstract magnitude from digits than it is from dice, presumably reflecting the relative frequency and familiarity of the stimuli. Alternatively, numerical formats could activate magnitude information in a *shared* but not necessarily abstract format. The delay for accessing magnitude information when presented with dice would then be attributed to the time it takes to convert the dice into the shared representation, potentially of digits. Disentangling these two alternatives is difficult. The current data show that there is sufficient similarity in processing of digits and dice to allow for cross-generalisation, but we cannot tell whether this is a different representation from either notation directly. We are hence cautious with the term *abstract* here and interpret the current data as evidence for a *shared* representation of magnitude for digit and dice stimuli. This interpretation allows for both explanations, an abstract representation or a representation in one numerical format only.

We have to be cautious when interpreting our data as it is hard to infer the source of the decodable signal (Coltheart, 2006; Henson, 2005). Our results show that it is possible to represent magnitude in a format-independent fashion but we cannot be certain whether this format-independent representation is *necessary* for normal number processing (Seron & Fias, 2006). It is, for example, possible that the task we asked participants to do resulted in the format-independent magnitude effect. Participants completed a 1-back task on magnitude which required them to think of the stimuli as representing magnitude. In future studies, it may be interesting to see whether magnitude can be decoded even if the task is completely orthogonal to magnitude processing, demonstrating whether attention to magnitude is a crucial aspect for such apparent shared representation.

Another caveat relates to the potential for covert semantic labelling to contribute to the magnitude effects. The Label Model was designed to account for task effects related to this. There was no time point at which this model correlated significantly with the data (Figure 7), but this is a null effect, and so we must be cautious in our inference. As there was no significant difference between the correlations of the data with the Magnitude Model and the Label Model, we cannot rule out a contribution of semantic labelling. However, the observation that the Magnitude Model provided a significant account of the data suggests that the ordinal structure in the model provided explanatory power, whereas we have no such clear information regarding the Label Model.

With our analysis, we are able to distinguish purely visual from higher-cognitive magnitude effects. The visual effects were much stronger and easier to decode than anything related to magnitude across our analyses. This is not surprising given the reliability of low-level visual signals, the time-locked nature of such signals to the stimulus, and the greater variability in individual processing times (even on a trial to trial basis) of higher-level cognitive functions. Looking at these visual effects, we also showed that dice produce a much stronger and clearer visual signal than digits. This again is not surprising given the visual dissimilarity within the non-symbolic stimuli such as dice: total luminance, for example, is lower for larger magnitudes than for smaller ones. In comparison, for digits the amount of visual information is relatively consistent across stimuli. This corresponds to results from previous fMRI MVPA studies consistently showing that non-symbolic stimuli (e.g., dots) resulted in higher magnitude decoding accuracy across the whole brain (Bulthé et al., 2014) and in the parietal lobes (Damarla & Just, 2013; Eger et al., 2009) than symbolic stimuli (i.e., digits). Bulthé et al. (2014) and Eger et al. (2009) controlled for some low-level visual features of the non-symbolic displays such as individual dot size, space between dots, total luminance, and total area of the stimuli. While controlling for these features limits the problem of visual dissimilarity across stimulus classes, some visual differences remain. For example, symbolic stimuli always consist of one item on the screen while non-symbolic stimuli consist of multiple items. These visual differences between stimulus classes may have led to higher decoding accuracy for dice than for digits in our study and previous studies. Our main results cannot be driven by such inevitable visual differences, as the key comparisons we make are based on a comparison across two different notations.

To our knowledge, the current study is the first to take a time-series decoding approach in the field of numerical cognition. Our results show that current analysis tools of MEG decoding are sensitive enough to distinguish between magnitudes. These methods offer many future avenues for the field of numerical cognition, as well as providing proof-of-concept that the methods can be applied to higher-level cognitive processes.

In summary, the results of the current study suggest that there is a shared magnitude representation regardless of symbolic notation. We also showed that there is a time shift in processing magnitude of different symbolic numerical formats with digits being accessed slightly earlier than dice. Although within-format classification is driven strongly by visual effects, we found that magnitude information across numerical formats can be decoded at a later stage in processing. By showing that magnitude is decodable, our study highlights that applying decoding to time-series data can be a useful approach for the field of numerical cognition.

## ACKNOWLEDGEMENTS

This research was supported by the ARC Centre of Excellence in Cognition and its Disorders, an ARC Future Fellowship (FT120100816) and ARC Discovery project (DP160101300) awarded to Thomas Carlson. A. Lina Teichmann and Tijl Grootswagers were supported by International Macquarie University Research Training Program Scholarships.

## Footnotes

1 Note, for research on the distinction between magnitude processing when accessed via symbolic and non-symbolic notations see e.g., Bulthé, De Smedt, … Op de Beeck (2014), Fias, Lammertyn, Reynvoet, Dupont, … Orban (2003), Libertus, Woldorff, … Brannon (2007), Piazza, Pinel, Le Bihan, … Dehaene (2007).

